## Supplementary Data for "Deep mutational scanning identifies variants of Cas1 and Cas2 that increase spacer acquisition in type II-A CRISPR-Cas systems"

### Supplementary Data 1. Heatmaps of global differential selection across Cas1, Cas2 and Csn2.

**A**

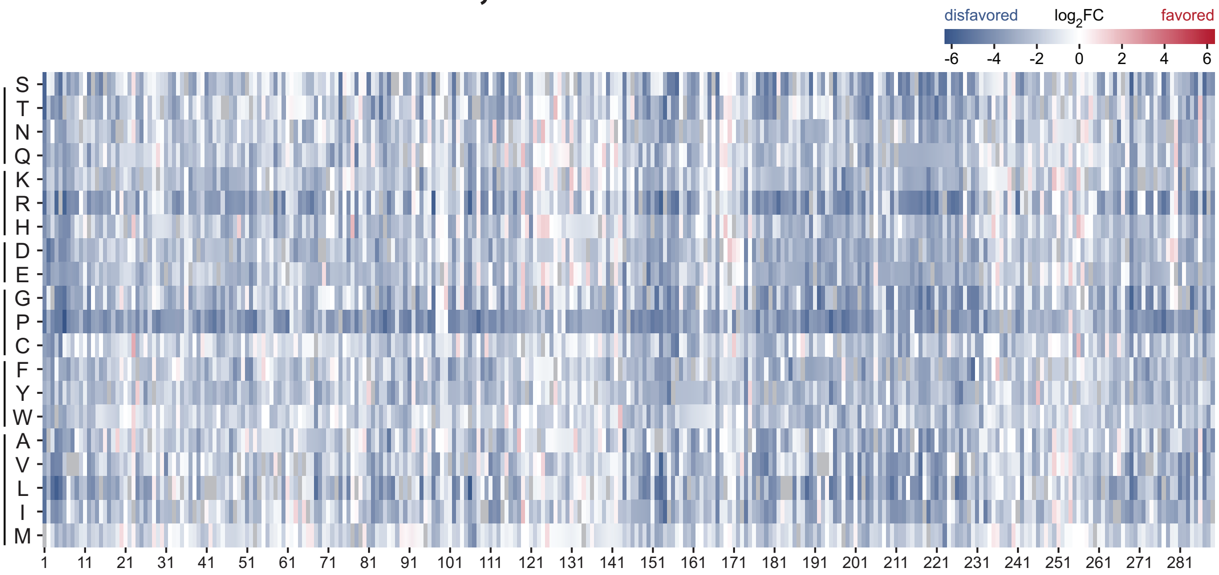


**B**

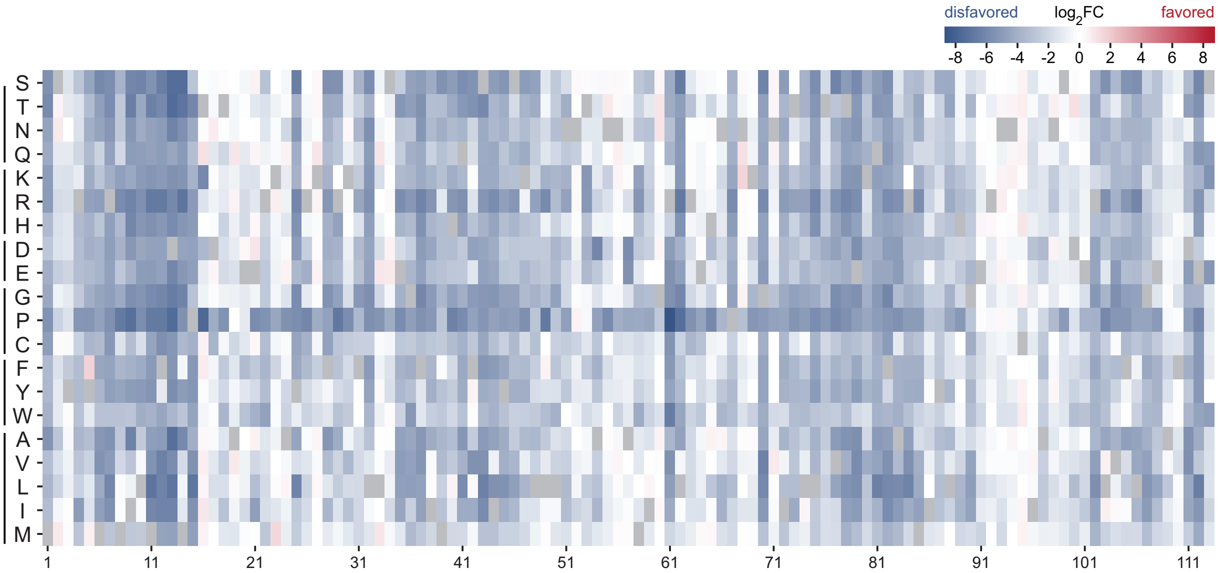


**C**

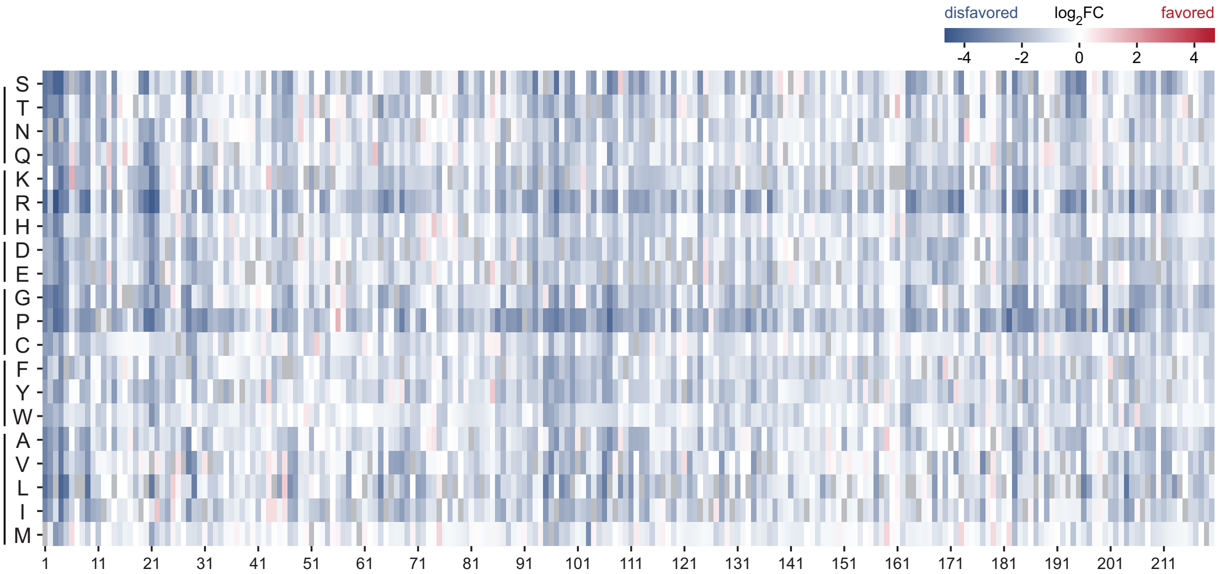


Global differential selection of amino acids across Cas1 (**A**), Cas2 (**B**) and Csn2 (**C**) according to DMS. Columns along the x-axis represent the positions in the protein sequence, and rows along the y-axis show the differential selection of each amino acid at that positions, colored by $log_2$ fold change ($log_2FC$).

### Supplementary Data 2. Positive differential selection and amino acid preferences of Cas1.

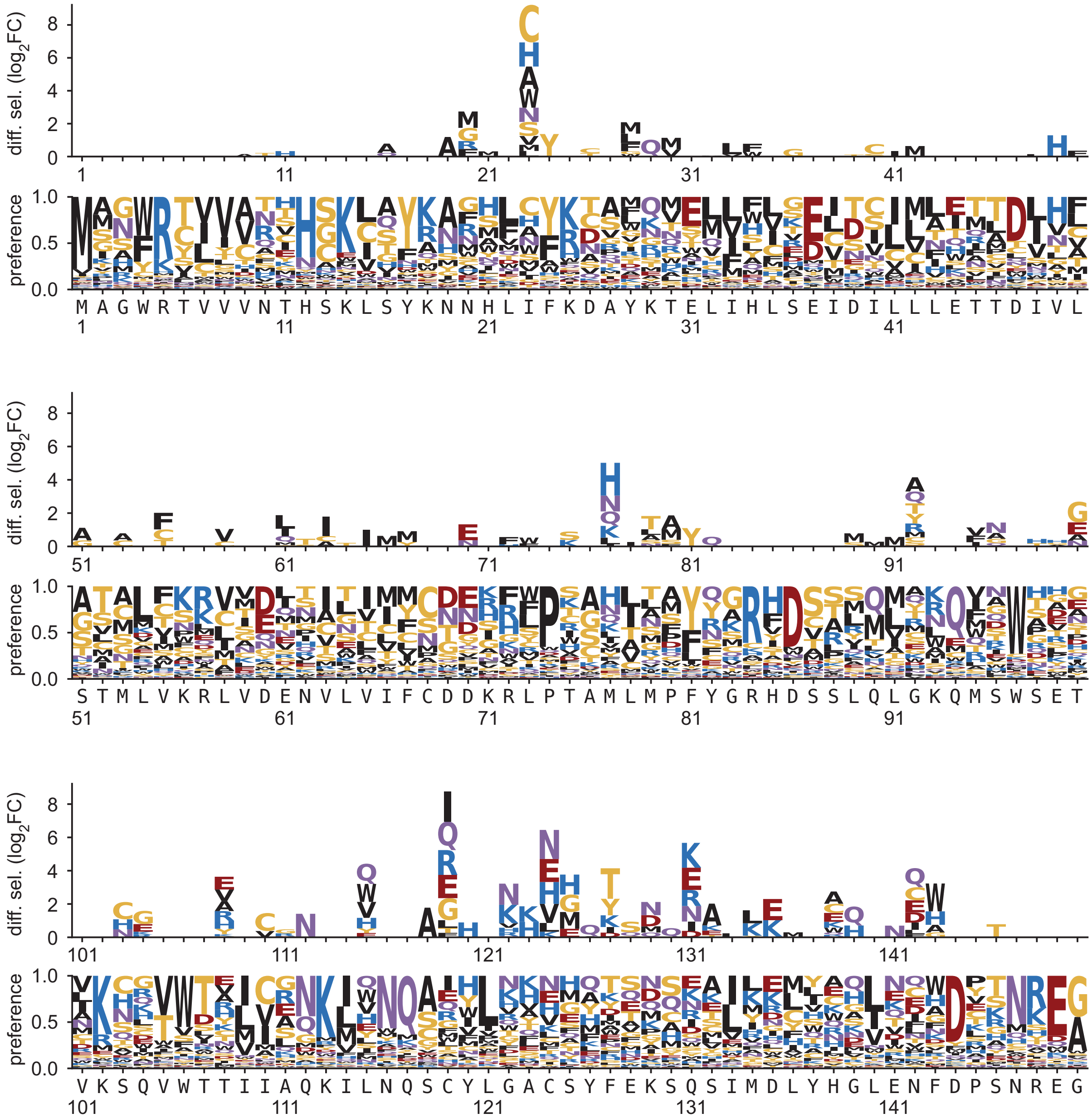


Positive differential selection (top) and amino acid preference (bottom) of Cas1 as determined by DMS. Figure continues to next page.

### Supplementary Data 2. Continued.

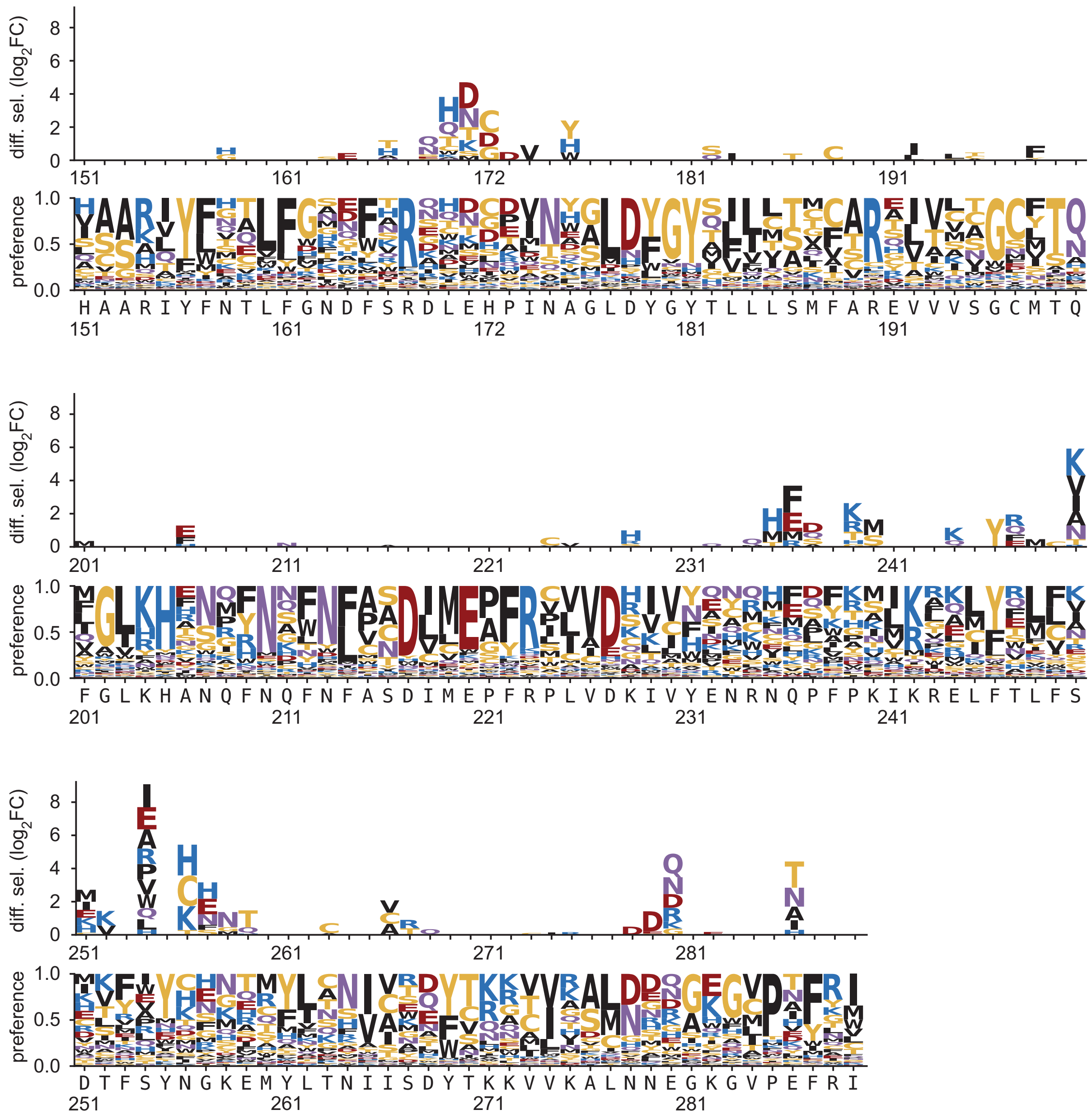

### Supplementary Data 3. Positive differential selection and amino acid preferences of Cas2.

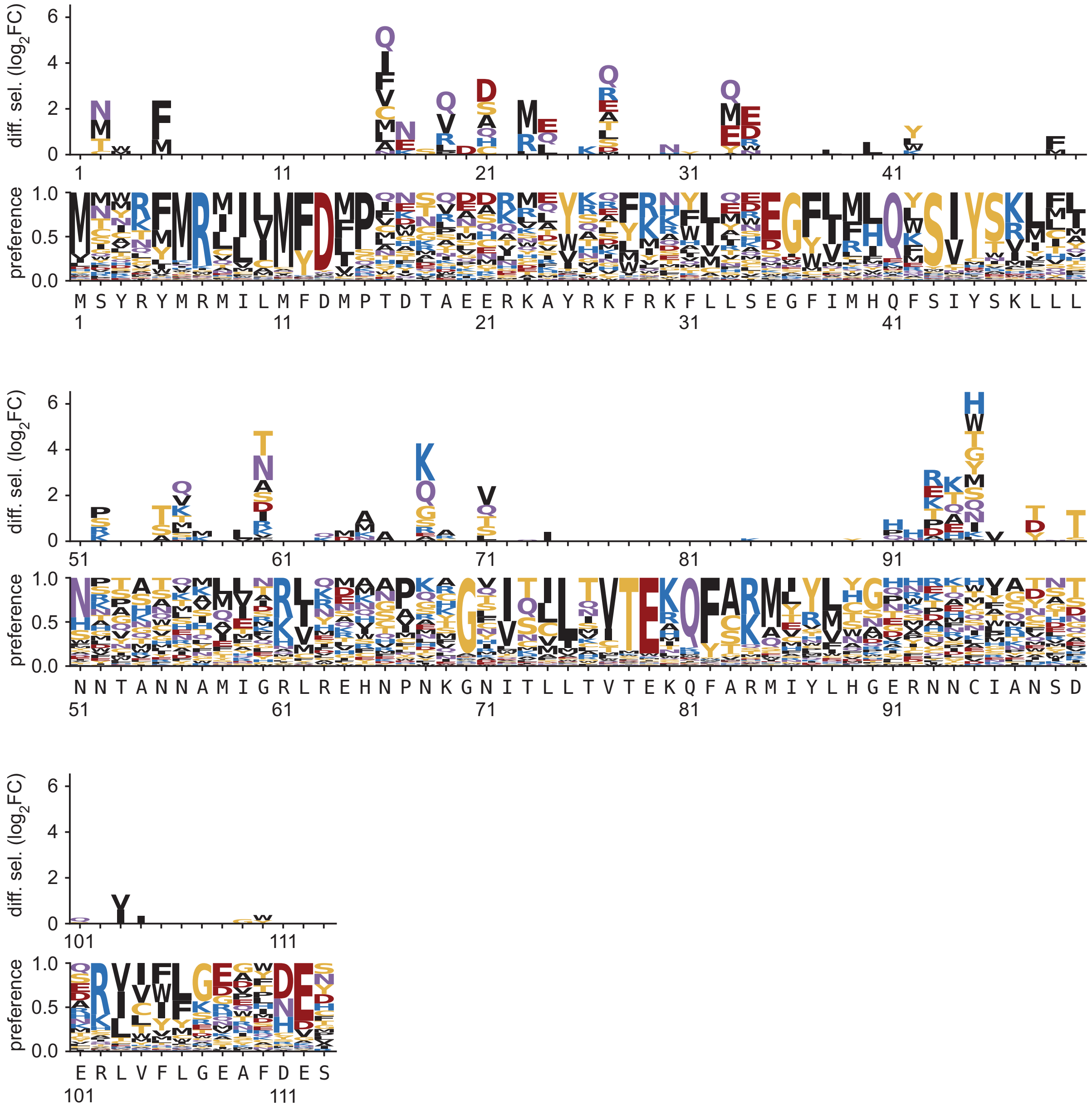


Positive differential selection (top) and amino acid preference (bottom) of Cas2 as determined by DMS.

### Supplementary Data 4. Positive differential selection and amino acid preferences of Csn2.

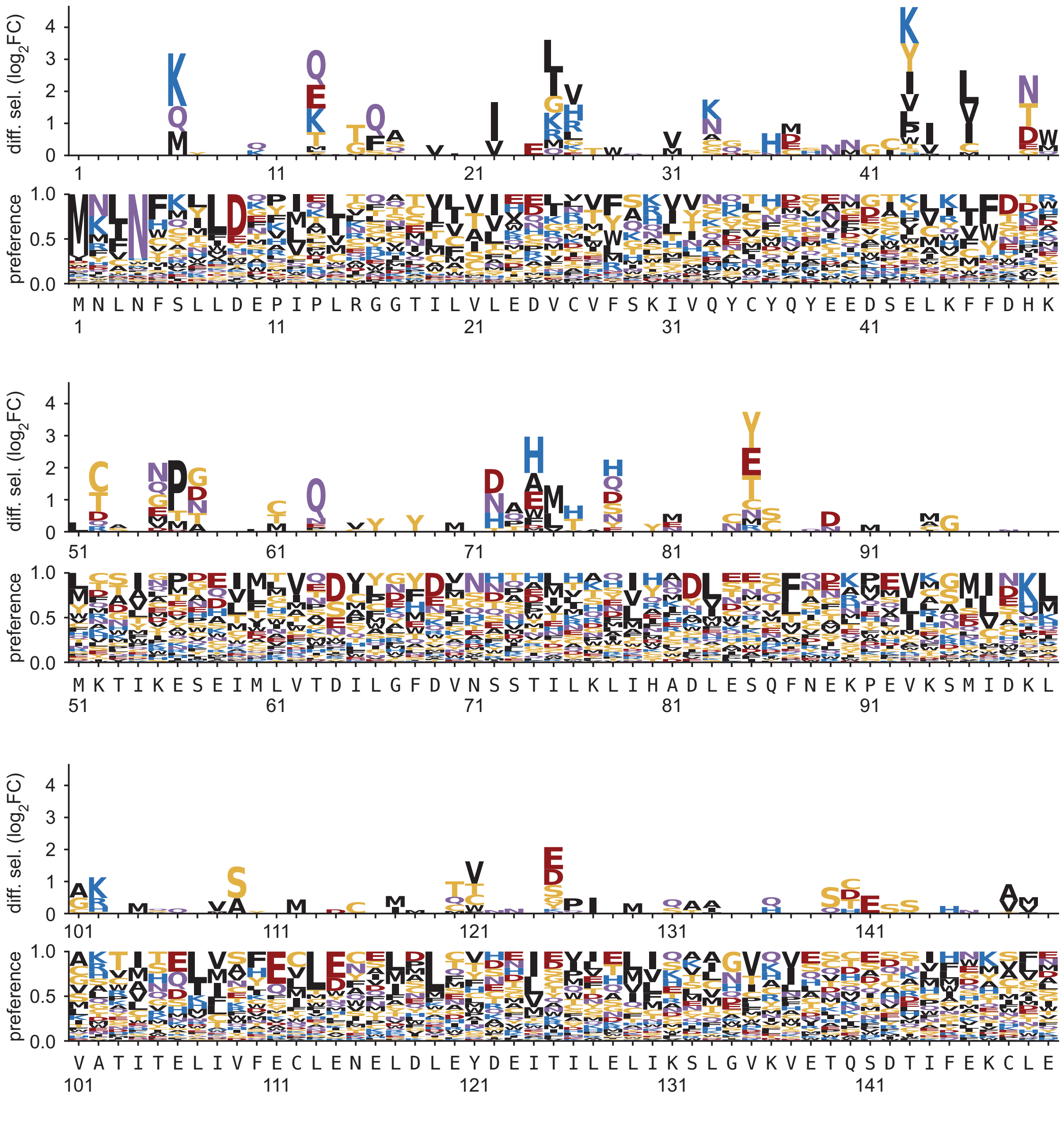


Positive differential selection (top) and amino acid preference (bottom) of Csn2 as determined by DMS. Figure continues to next page.

### Supplementary Data 4. Continued.

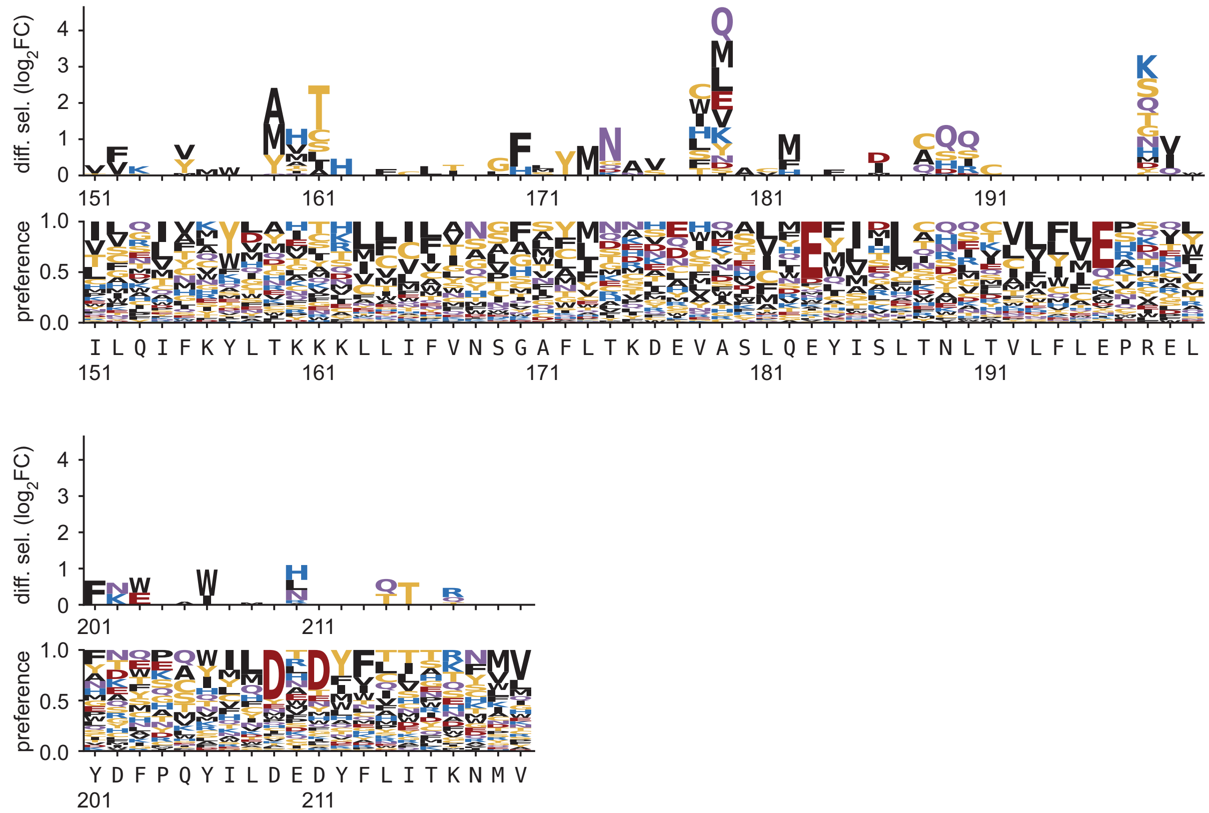

### Supplementary Data 5. Positive and negative differential selection of Cas1.

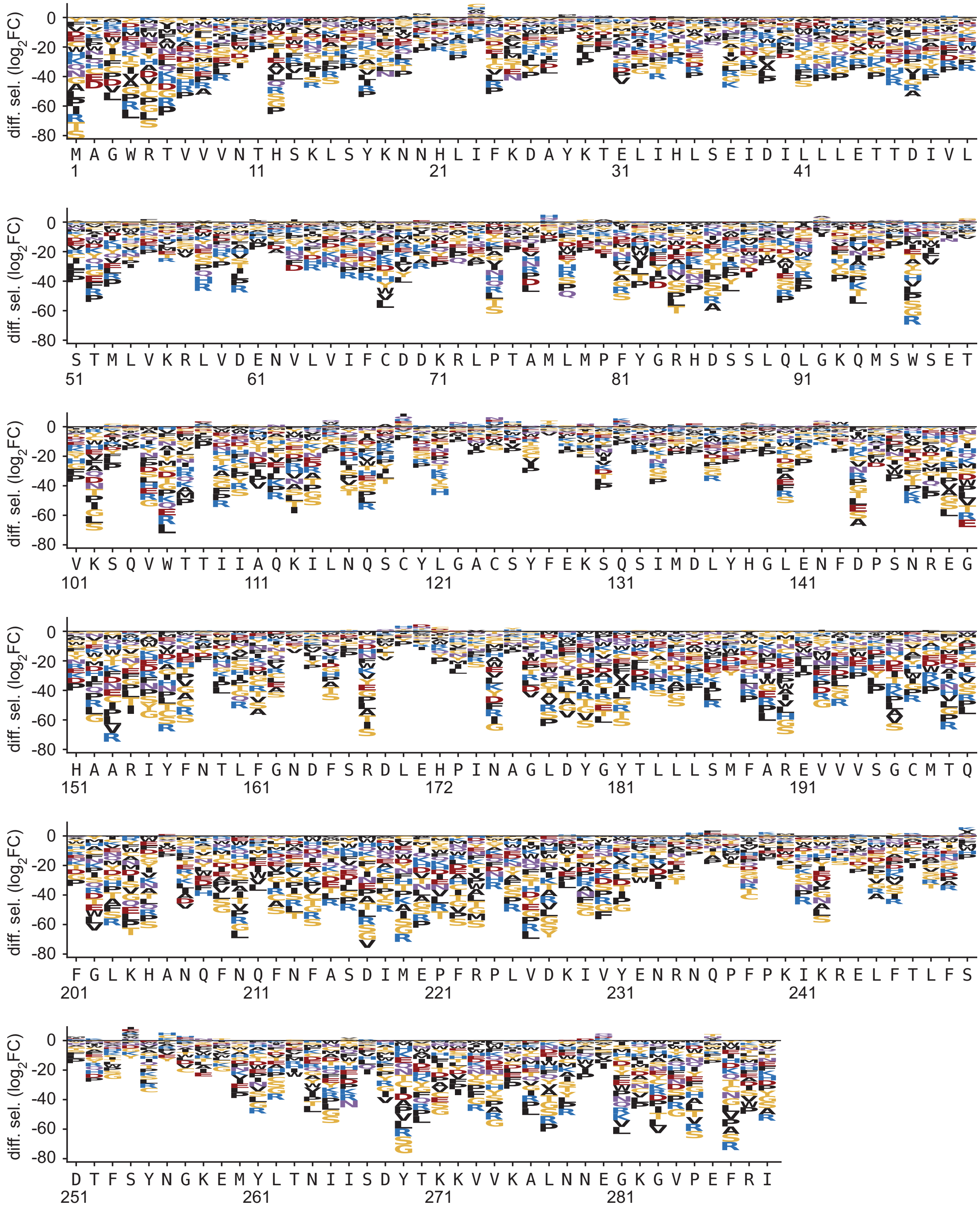

### Supplementary Data 6. Positive and negative differential selection of Cas2.

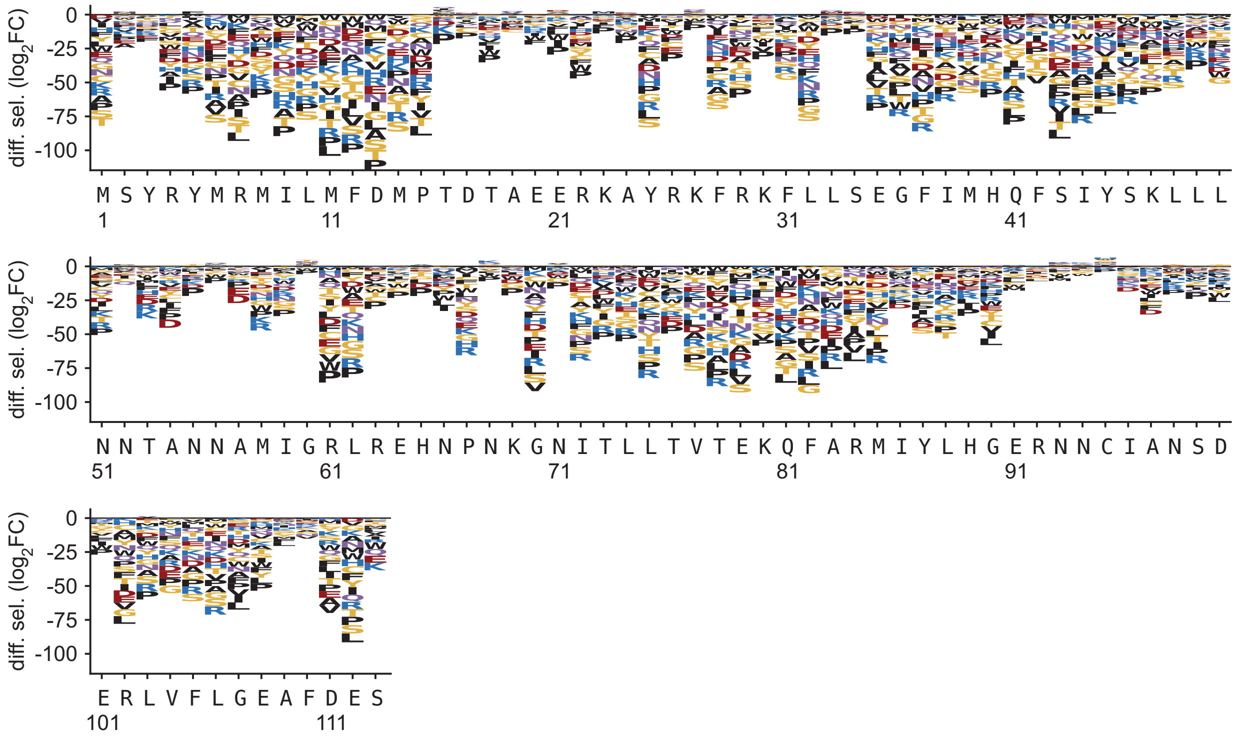

### Supplementary Data 7. Positive and negative differential selection of Csn2.

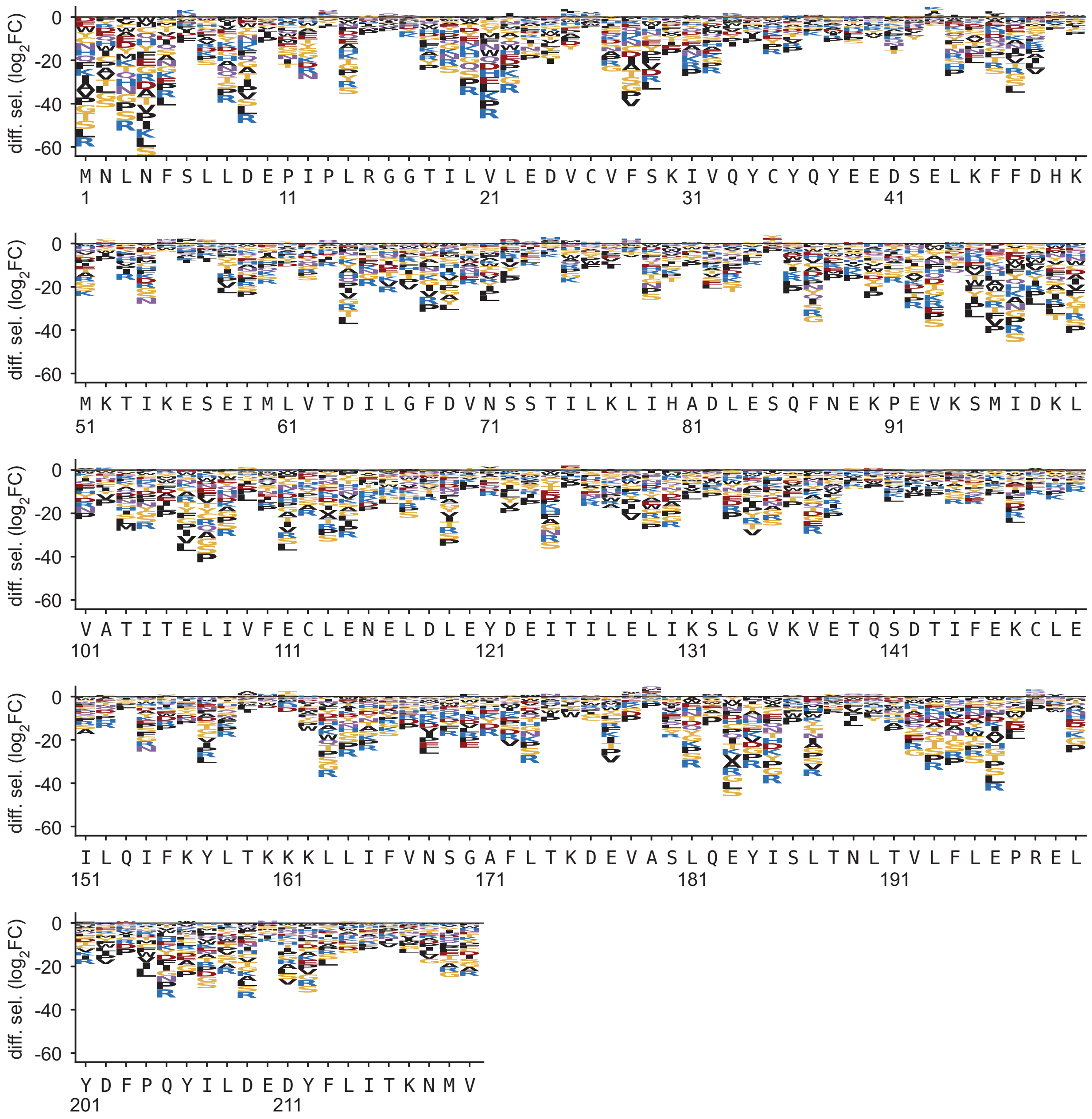

### Supplementary Data 8. Alignment of Cas1 protein sequences.

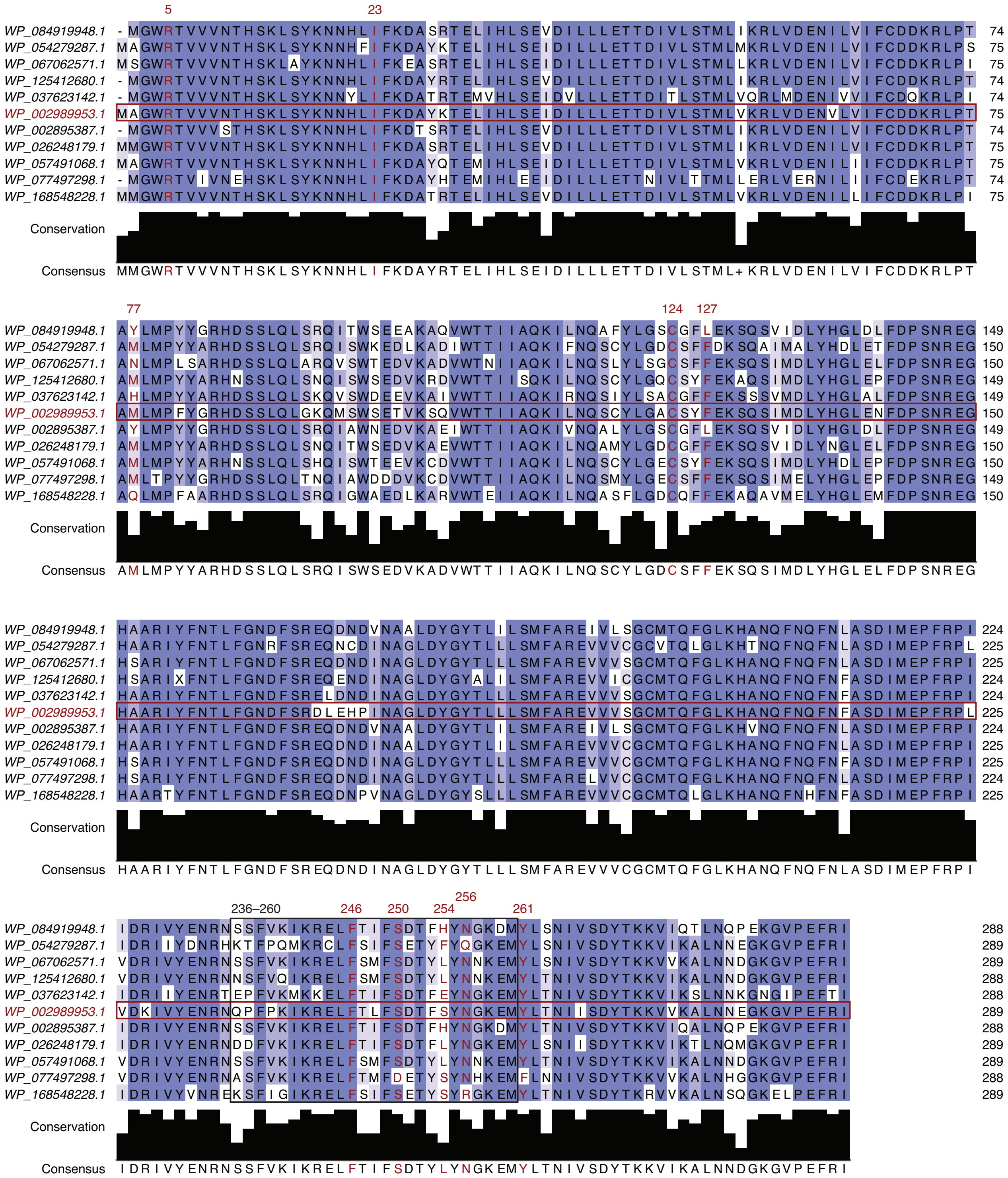


Alignment of Cas1 protein sequences, colored by sequence identity. The sequence of *S. pyogenes* Cas1 is highlighted with a red box, select positions are highlighted in red font, and residues 236–260 are highlighted with a black box.

### Supplementary Data 9. Alignment of Cas2 protein sequences.

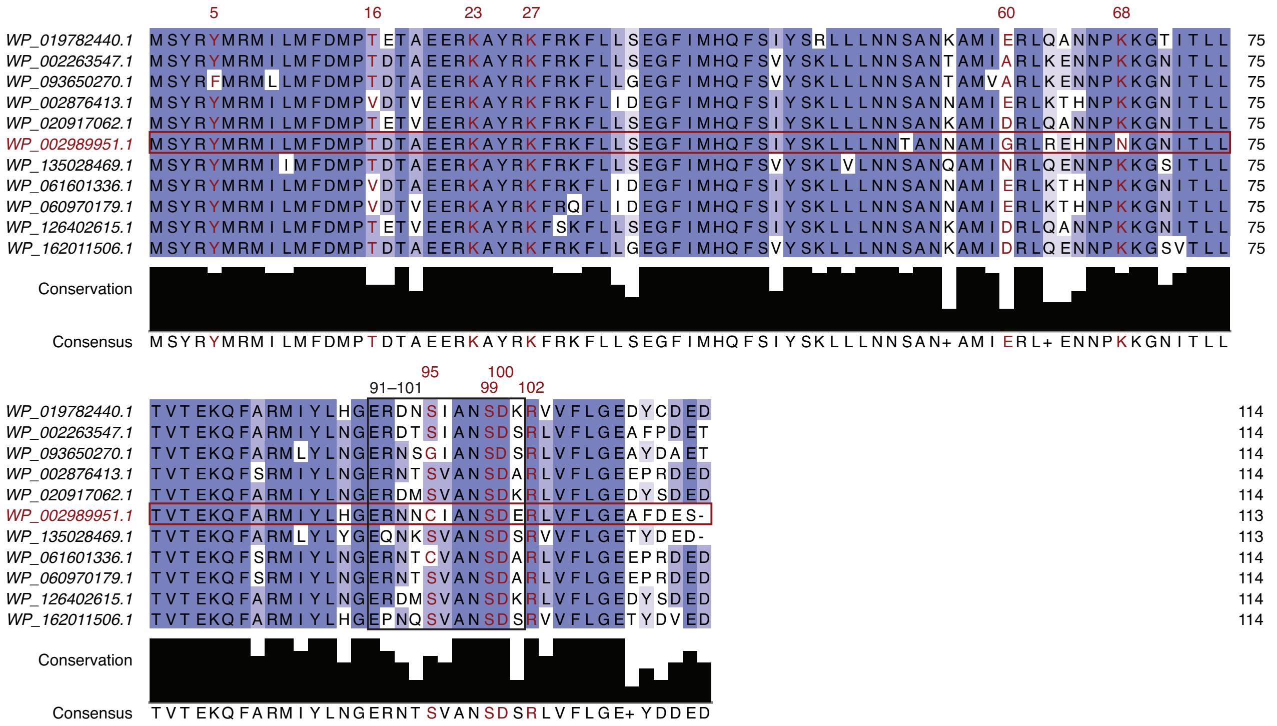


Alignment of Cas2 protein sequences, colored by sequence identity. The sequence of *S. pyogenes* Cas2 is highlighted with a red box, select positions are highlighted in red font, and residues 91–101 are highlighted with a black box.

### Supplementary Data 10. Alignment of Csn2 protein sequences.

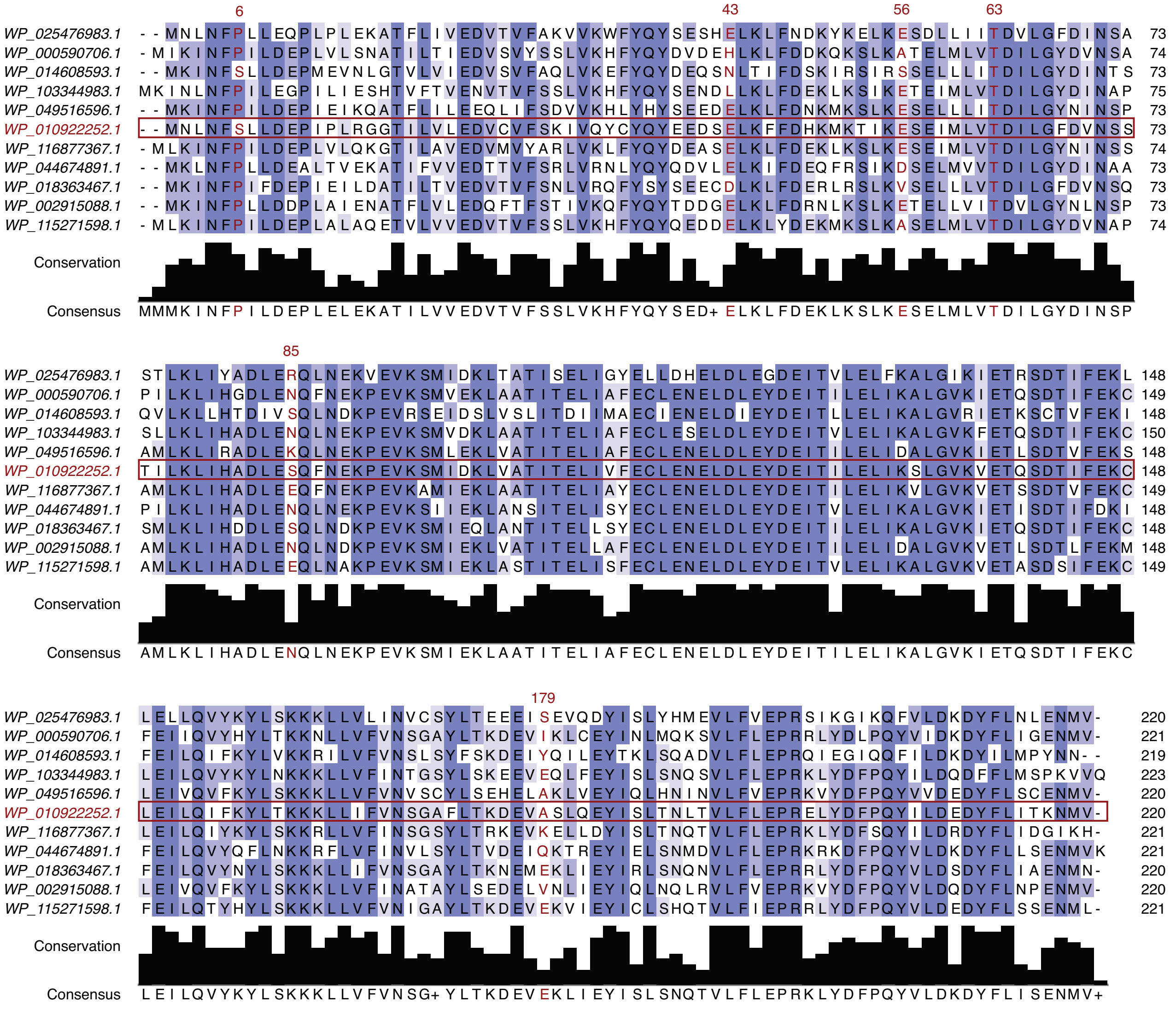


Alignment of Csn2 protein sequences, colored by sequence identity. The sequence of *S. pyogenes* Csn2 is highlighted with a red box and select positions are highlighted in red font.
