## Supplementary Tables for "Deep mutational scanning identifies variants of Cas1 and Cas2 that increase spacer acquisition in type II-A CRISPR-Cas systems"

**Supplementary Table S1.** Plasmids used in this study.

| Plasmid | Description | Source | Construction notes* |
|---|---|---|---|
| pCH2 | *S. pyogenes* type II-A CRISPR-Cas system (Δ*tracr-L*) with a single repeat and mNG reporter in pC194-based vector | This study | Gibson assembly: oCH8+oCH7 (pRAH13), oCH5+oCH6 (mNG) |
| pCH4 | *S. pyogenes* type II-A CRISPR-Cas system (Δ*tracr-L*) with *spc1(ATG)* and mNG reporter in pC194-based vector | This study | Gibson assembly: oCH8+oCH7 (pRAH11), oCH5+oCH6 (mNG) |
| pCH6 | *S. pyogenes* type II-A CRISPR-Cas system (Δ*tracr-L*) with *spc1(TAG)* and mNG reporter in pC194-based vector | This study | Gibson assembly: oCH8+oCH7 (pRAH16), oCH5+oCH6 (mNG) |
| pCH7 | Like pCH2, with *d* variant | This study | Gibson assembly: oCH21+oRAH102 (pCH2), oCH22+oRAH91 (pCH2) |
| pCH8 | Like pCH4, with *d* variant | This study | Gibson assembly: oCH21+oRAH102 (pCH4), oCH22+oRAH91 (pCH4) |
| pCH9 | Like pCH6, with *d* variant | This study | Gibson assembly: oCH21+oRAH102 (pCH6), oCH22+oRAH91 (pCH6) |
| pCH10 | Like pCH2, with *cs* variant | This study | Gibson assembly: oCH23+oRAH102 (pCH2), oCH24+oRAH91 (pCH2) |
| pCH11 | Like pCH4, with *cs* variant | This study | Gibson assembly: oCH23+oRAH102 (pCH4), oCH24+oRAH91 (pCH4) |
| pCH12 | Like pCH6, with *cs* variant | This study | Gibson assembly: oCH23+oRAH102 (pCH6), oCH24+oRAH91 (pCH6) |
| pCH13 | Like pCH2, with *p* variant | This study | Gibson assembly: oCH25+oRAH102 (pCH2), oCH26+oRAH91 (pCH2) |
| pCH14 | Like pCH4, with *p* variant | This study | Gibson assembly: oCH25+oRAH102 (pCH4), oCH26+oRAH91 (pCH4) |
| pCH15 | Like pCH6, with *p* variant | This study | Gibson assembly: oCH25+oRAH102 (pCH6), oCH26+oRAH91 (pCH6) |

| | | | |
|---|---|---|---|
| pCH16 | Like pCH2, with sgRNA variant | This study | Gibson assembly:<br>oCH27+oRAH102 (pCH2), oCH28+oRAH91 (pCH2), oCH31 stitching primer |
| pCH17 | Like pCH4, with sgRNA variant | This study | Gibson assembly:<br>oCH27+oRAH102 (pCH4), oCH28+oRAH91 (pCH4), oCH31 stitching primer |
| pCH18 | Like pCH6, with sgRNA variant | This study | Gibson assembly:<br>oCH27+oRAH102 (pCH6), oCH28+oRAH91 (pCH6), oCH31 stitching primer |
| pCH19 | Like pCH2, with *p+d* variant | This study | Gibson assembly:<br>oCH29+oRAH102 (pCH2), oCH30+oRAH91 (pCH2) |
| pCH20 | Like pCH4, with *p+d* variant | This study | Gibson assembly:<br>oCH29+oRAH102 (pCH4), oCH30+oRAH91 (pCH4) |
| pCH21 | Like pCH6, with *p+d* variant | This study | Gibson assembly:<br>oCH29+oRAH102 (pCH6), oCH30+oRAH91 (pCH6) |
| pRAH11 | *S. pyogenes* type II-A CRISPR-Cas system (Δ*tracr-L*) with *spc1(ATG)* and msfGFP reporter in pC194-based vector | This study | Gibson assembly:<br>oRAH62+oRAH73 (pGG32-Δ*tracr-L*, Workman *et al.*[1]), oRAH74+oRAH70 (pGG32-Δ*tracr-L*), oRAH69+oRAH71 (msfGFP), oRAH72+oRAH61 (pGG32-Δ*tracr-L*) |
| pRAH13 | *S. pyogenes* type II-A CRISPR-Cas system (Δ*tracr-L*) with a single repeat and msfGFP reporter in pC194-based vector | This study | Gibson assembly:<br>oRAH62+oRAH70 (pGG32-Δ*tracr-L*, Workman *et al.*[1]), oRAH69+oRAH71 (msfGFP), oRAH72+oRAH61 (pGG32-Δ*tracr-L*) |
| pRAH16 | *S. pyogenes* type II-A CRISPR-Cas system (Δ*tracr-L*) with *spc1(TAG)* and msfGFP reporter in pC194-based vector | This study | Gibson assembly:<br>oRAH62+oRAH82 (pGG32-Δ*tracr-L*, Workman *et al.*[1]), oRAH81+oRAH70 (pGG32-Δ*tracr-L*), oRAH69+oRAH71 (msfGFP), oRAH72+oRAH61 (pGG32-Δ*tracr-L*) |
| pRAH49-1 | *S. pyogenes* type II-A CRISPR-Cas system (Δ*tracr-L* and *cs* variant) with a single repeat and ErmR reporter in pC194-based vector; also denoted *wt* and *cs* | This study | Gibson assembly:<br>oRAH320+oRAH97 (pCH10), oRAH86+oRAH318 (pCH10), oRAH319+oRAH321 (pE194, Horinouchi *et al.*[2]) |

| | | | |
|---|---|---|---|
| pRAH50 | Like pRAH49-1, with *spc1(ATG)*; also denoted *cs* + *spc1(ATG)* | This study | Gibson assembly:<br>oRAH320+oRAH97 (pCH11), oRAH86+oRAH318 (pCH11), oRAH319+oRAH321 (pE194, Horinouchi *et al.*[2]) |
| pRAH77 | Like pRAH49-1, with Δ*csn2*; also denoted *cs* + Δ*csn2* | This study | Blunt-end ligation:<br>oRAH433+oRAH434 (pRAH49-1) |
| pRAH88 | Like pRAH49-1, with Csn2 S6K | This study | Gibson assembly:<br>oRAH455+oRAH61 (pRAH49-1), oRAH62+oRAH454 (pRAH49-1) |
| pRAH89 | Like pRAH49-1, with Csn2 E56P | This study | Gibson assembly:<br>oRAH457+oRAH61 (pRAH49-1), oRAH62+oRAH456 (pRAH49-1) |
| pRAH90 | Like pRAH49-1, with Csn2 T63Q | This study | Gibson assembly:<br>oRAH458+oRAH61 (pRAH49-1), oRAH62+oRAH459 (pRAH49-1) |
| pRAH91 | Like pRAH49-1, with Csn2 A179Q | This study | Gibson assembly:<br>oRAH460+oRAH61 (pRAH49-1), oRAH62+oRAH461 (pRAH49-1) |
| pRAH92 | Like pRAH49-1, with Csn2 E43K | This study | Gibson assembly:<br>oRAH462+oRAH61 (pRAH49-1), oRAH62+oRAH463 (pRAH49-1) |
| pRAH93 | Like pRAH49-1, with Csn2 E43Y | This study | Gibson assembly:<br>oRAH464+oRAH61 (pRAH49-1), oRAH62+oRAH465 (pRAH49-1) |
| pRAH94 | Like pRAH49-1, with Csn2 S85E | This study | Gibson assembly:<br>oRAH466+oRAH61 (pRAH49-1), oRAH62+oRAH467 (pRAH49-1) |
| pRAH95 | Like pRAH49-1, with Csn2 S85Y | This study | Gibson assembly:<br>oRAH468+oRAH61 (pRAH49-1), oRAH62+oRAH469 (pRAH49-1) |
| pRAH158 | Like pRAH49-1, with Cas1 I23C | This study | Gibson assembly:<br>oRAH575+oRAH61 (pRAH49-1), oRAH62+oRAH576 (pRAH49-1) |
| pRAH159 | Like pRAH49-1, with Cas1 M77H | This study | Gibson assembly:<br>oRAH577+oRAH61 (pRAH49-1), oRAH62+oRAH578 (pRAH49-1) |
| pRAH162 | Like pRAH49-1, with Cas1 C124N | This study | Gibson assembly:<br>oRAH583+oRAH61 (pRAH49-1), oRAH62+oRAH584 (pRAH49-1) |
| pRAH163 | Like pRAH49-1, with Cas1 F127T | This study | Gibson assembly:<br>oRAH585+oRAH61 (pRAH49-1), oRAH62+oRAH586 (pRAH49-1) |

| | | | |
|---|---|---|---|
| pRAH164 | Like pRAH49-1, with Cas1 F246Y | This study | Gibson assembly: oRAH587+oRAH61 (pRAH49-1), oRAH62+oRAH588 (pRAH49-1) |
| pRAH165 | Like pRAH49-1, with Cas1 S250K | This study | Gibson assembly: oRAH589+oRAH61 (pRAH49-1), oRAH62+oRAH590 (pRAH49-1) |
| pRAH166 | Like pRAH49-1, with Cas1 S254I | This study | Gibson assembly: oRAH591+oRAH61 (pRAH49-1), oRAH62+oRAH592 (pRAH49-1) |
| pRAH167 | Like pRAH49-1, with Cas1 N256H | This study | Gibson assembly: oRAH593+oRAH61 (pRAH49-1), oRAH62+oRAH594 (pRAH49-1) |
| pRAH168 | Like pRAH49-1, with Cas2 Y5F | This study | Gibson assembly: oRAH595+oRAH61 (pRAH49-1), oRAH62+oRAH596 (pRAH49-1) |
| pRAH169 | Like pRAH49-1, with Cas2 T16Q | This study | Gibson assembly: oRAH597+oRAH61 (pRAH49-1), oRAH62+oRAH598 (pRAH49-1) |
| pRAH170 | Like pRAH49-1, with Cas2 K23M | This study | Gibson assembly: oRAH599+oRAH61 (pRAH49-1), oRAH62+oRAH600 (pRAH49-1) |
| pRAH171 | Like pRAH49-1, with Cas2 K27Q | This study | Gibson assembly: oRAH601+oRAH61 (pRAH49-1), oRAH62+oRAH602 (pRAH49-1) |
| pRAH172 | Like pRAH49-1, with Cas2 G60T | This study | Gibson assembly: oRAH603+oRAH61 (pRAH49-1), oRAH62+oRAH604 (pRAH49-1) |
| pRAH173 | Like pRAH49-1, with Cas2 N68K | This study | Gibson assembly: oRAH605+oRAH61 (pRAH49-1), oRAH62+oRAH606 (pRAH49-1) |
| pRAH174 | Like pRAH49-1, with Cas2 C95H | This study | Gibson assembly: oRAH607+oRAH61 (pRAH49-1), oRAH62+oRAH608 (pRAH49-1) |
| pRAH175 | Like pRAH49-1, with Cas2 D100T | This study | Gibson assembly: oRAH609+oRAH61 (pRAH49-1), oRAH62+oRAH610 (pRAH49-1) |
| pRAH185 | Like pRAH49-1, with Cas1 M77H, Cas2 T16Q | This study | Gibson assembly: oRAH61+oRAH309 (pRAH159), oRAH62+oRAH307 (pRAH159), oRAH306+oRAH308 (pRAH169) |
| pRAH186 | Like pRAH49-1, with Cas1 M77H, Csn2 S85E | This study | Gibson assembly: oRAH61+oRAH313 (pRAH159), oRAH62+oRAH308 (pRAH159), oRAH309+oRAH312 (pRAH94) |

| | | | |
|---|---|---|---|
| pRAH187 | Like pRAH49-1, with Cas2 T16Q, Csn2 S85E | This study | Gibson assembly:<br>oRAH62+oRAH308 (pRAH169), oRAH61+oRAH313 (pRAH169), oRAH309+oRAH312 (pRAH94) |
| pRAH188 | Like pRAH49-1, with Cas1 M77H, Cas2 T16Q, Csn2 S85E | This study | Gibson assembly:<br>oRAH61+oRAH313 (pRAH159), oRAH62+oRAH307 (pRAH159), oRAH306+oRAH308 (pRAH169), oRAH309+oRAH312 (pRAH94) |
| pRAH189 | Like pRAH49-1, with Cas2 Y5F T16Q | This study | Gibson assembly:<br>oRAH595+oRAH61 (pRAH169), oRAH62+oRAH596 (pRAH169) |
| pRAH201 | Like pRAH49-1, with Cas1 M77H, Cas2 Y5F | This study | Gibson assembly:<br>oRAH61+oRAH309 (pRAH159), oRAH62+oRAH307 (pRAH159), oRAH306+oRAH308 (pRAH168) |
| pRAH202 | Like pRAH49-1, with Cas1 M77H, Cas2 Y5F T16Q | This study | Gibson assembly:<br>oRAH61+oRAH309 (pRAH159), oRAH62+oRAH307 (pRAH159), oRAH306+oRAH308 (pRAH189) |
| pRAH216 | *S. pyogenes* type II-A CRISPR-Cas system with a single repeat in pLZ12-based vector | This study | Gibson assembly:<br>oRAH83+oRAH389 (pGG32, Heler *et al.*[3]), oRAH390+oRAH84 (pPN75) |
| pRAH218 | Like pRAH216, with Cas2 T16Q | This study | Gibson assembly:<br>oRAH83+oRAH389 (pRAH169), oRAH390+oRAH84 (pPN75) |
| pRAH219 | Like pRAH216, with Cas2 Y5F T16Q | This study | Gibson assembly:<br>oRAH83+oRAH389 (pRAH189), oRAH390+oRAH84 (pPN75) |
| pRAH220 | Like pRAH216, with Cas1 M77H, Cas2 Y5F T16Q | This study | Gibson assembly:<br>oRAH83+oRAH389 (pRAH202), oRAH390+oRAH84 (pPN75) |
| pRAH225 | *S. pyogenes* type II-A Δ*cas1* Δ*cas2* Δ*csn2* CRISPR-Cas system (Δ*tracr-L*) with a single repeat in pLZ12-based vector; also denoted Δ*cas1/cas2/csn2* | This study | Blunt-end ligation:<br>oRAH434+oRAH526 (pRAH226) |
| pRAH226 | *S. pyogenes* type II-A CRISPR-Cas system (Δ*tracr-L*) with a single repeat in pLZ12-based vector; also denoted wt and pWT | This study | Gibson assembly:<br>oHK55+oRAH299 (pGG32-Δ*tracr-L*, Workman *et al.*[1]), oRAH298+oRAH389 (pRAH216), oRAH390+oHK156 (pRAH216) |
| pRAH227 | Like pRAH226, with Cas2 T16Q; also denoted variant 1 and pVARIANT1 | This study | Gibson assembly:<br>oHK55+oRAH299 (pGG32-Δ*tracr-L*, Workman *et al.*[1]), oRAH298+oRAH389 (pRAH218), oRAH390+oHK156 (pRAH216) |

| | | | |
|---|---|---|---|
| pRAH228 | Like pRAH226, with Cas2 Y5F T16Q; also denoted variant 2 and pVARIANT2 | This study | Gibson assembly:<br>oHK55+oRAH299 (pGG32-Δ*tracr-L*, Workman *et al.*[1]), oRAH298+oRAH389 (pRAH219), oRAH390+oHK156 (pRAH216) |
| pRAH229 | Like pRAH226, with Cas1 M77H, Cas2 Y5F T16Q; also denoted variant 3 and pVARIANT3 | This study | Gibson assembly:<br>oHK55+oRAH299 (pGG32-Δ*tracr-L*, Workman *et al.*[1]), oRAH298+oRAH389 (pRAH220), oRAH390+oHK156 (pRAH216) |
| pRAH230 | Like pRAH49-1, with Cas1 Y261A | This study | Gibson assembly:<br>oRAH83+oRAH680 (pRAH49-1), oRAH679+oRAH84 (pRAH49-1) |
| pRAH231 | Like pRAH49-1, with Cas2 R102A | This study | Gibson assembly:<br>oRAH83+oRAH682 (pRAH49-1), oRAH681+oRAH84 (pRAH49-1) |
| pRAH232 | Like pRAH49-1, with Cas1 Y261A, Cas2 R102A | This study | Gibson assembly:<br>oRAH83+oRAH680 (pRAH49-1), oRAH679+oRAH682 (pRAH49-1), oRAH681+oRAH84 (pRAH49-1) |
| pRAH233 | Like pRAH49-1, with Cas1 Y261R, Cas2 R102Y | This study | Gibson assembly:<br>oRAH83+oRAH684 (pRAH49-1), oRAH683+oRAH686 (pRAH49-1), oRAH685+oRAH84 (pRAH49-1) |
| pRAH234 | Like pRAH49-1, with Cas1 Y261F | This study | Gibson assembly:<br>oRAH83+oRAH688 (pRAH49-1), oRAH687+oRAH84 (pRAH49-1) |
| pRAH235 | Like pRAH49-1, with Cas2 R102K | This study | Gibson assembly:<br>oRAH83+oRAH690 (pRAH49-1), oRAH689+oRAH84 (pRAH49-1) |
| pRAH236 | Like pRAH49-1, with Cas1 R5A | This study | Gibson assembly:<br>oRAH83+oRAH692 (pRAH49-1), oRAH691+oRAH84 (pRAH49-1) |
| pRAH237 | Like pRAH49-1, with Cas1 R5K | This study | Gibson assembly:<br>oRAH83+oRAH694 (pRAH49-1), oRAH693+oRAH84 (pRAH49-1) |
| pRAH238 | Like pRAH49-1, with Cas2 S99A | This study | Gibson assembly:<br>oRAH83+oRAH696 (pRAH49-1), oRAH695+oRAH84 (pRAH49-1) |
| pRAH241 | Like pRAH49-1, with Cas1 H205A E220A; also denoted dCas1 | This study | Gibson assembly:<br>oRAH83+oRAH698 (pRAH49-1), oRAH699+oRAH84 (pRAH49-1), oRAH697 and oRAH700 stitching primers |
| pRAH243 | Like pRAH49-1, with Cas2 R102Y | This study | Gibson assembly:<br>oRAH83+oRAH686 (pRAH49-1), oRAH685+oRAH84 (pRAH49-1) |

| | | | |
|---|---|---|---|
| pRAH244 | Like pRAH49-1, with Cas1 Y261R | This study | Gibson assembly:<br>oRAH83+oRAH684 (pRAH49-1), oRAH683+oRAH84 (pRAH49-1) |
| pRAH246 | Hexahistidyl-tagged SUMO fusion of *S. pyogenes* Cas1 in pET28-based vector | This study | Gibson assembly:<br>oRAH702+oRAH611 (Addgene #112793), oRAH701+oRAH703 (pRAH49-1) |
| pRAH247 | Hexahistidyl-Twin-Strep-II-tagged SUMO fusion of *S. pyogenes* Cas2 in pET28-based vector | This study | Gibson assembly:<br>oRAH612+oRAH611 (Addgene #112793), oRAH702+oRAH634 (Addgene#112794), oRAH704+oRAH705 (pRAH49-1) |
| pRAH248 | Like pRAH246, with Cas1 R5A | This study | Gibson assembly:<br>oRAH702+oRAH611 (Addgene #112793), oRAH701+oRAH706 (pRAH236) |
| pRAH249 | Like pRAH246, with Cas1 R5K | This study | Gibson assembly:<br>oRAH702+oRAH611 (Addgene #112793), oRAH701+oRAH707 (pRAH49-1) |

* Templates are listed in brackets.

**Supplementary Table S2.** Oligonucleotide primers used in this study.

| Primer | Sequence |
|---|---|
| Rnd2forUniv | aatgatacggcgaccaccgagatctacactctttccctacacgacgctcttcc |
| cym-TruSeq1 | caagcagaagacggcatacgagatCGTGATgtgactggagttcagacgtgtgctcttcc |
| cym-TruSeq2 | caagcagaagacggcatacgagatACATCGgtgactggagttcagacgtgtgctcttcc |
| cym-TruSeq3 | caagcagaagacggcatacgagatGCCTAAgtgactggagttcagacgtgtgctcttcc |
| cym-TruSeq5 | caagcagaagacggcatacgagatCACTGTgtgactggagttcagacgtgtgctcttcc |
| cym-TruSeq8 | caagcagaagacggcatacgagatTCAAGTgtgactggagttcagacgtgtgctcttcc |
| cym-TruSeq9 | caagcagaagacggcatacgagatCTGATCgtgactggagttcagacgtgtgctcttcc |
| cym-TruSeq10 | caagcagaagacggcatacgagatAAGCTAgtgactggagttcagacgtgtgctcttcc |
| oCH5 | cgatgataacgtgagcaaaggtgaagaggataatatgg |
| oCH6 | ctttctcaactattatttgtacagctcatccatacccatc |
| oCH7 | cacctttgctcacgttatcatcggcaatgttgttttggg |
| oCH8 | ggatgagctgtacaaataatagttgagaaagagggttaataccagcag |
| oCH9 | gttgtttcaggtcacaaagctttaattg |
| oCH10 | ttttaaaaatatctcttgcccccttctggcaagttgggatgtttatc |
| oCH11 | ggcaagagatatttttaaaacaatgaatagg |
| oCH12 | ctgacgaaagtccaagggtttattg |
| oCH13 | aacccttggactttcgtcagagggaatttaacaaaaatgcgtgccac |
| oCH14 | gaccctgaatatctttcaattgctc |
| oCH21 | tttaacttgtcatgctgttttgaatggttccaacaag |
| oCH22 | ccattcaaaacagcatgacaagttaaaataaggctagtccgttatcaac |
| oCH23 | gctatgctgtttctaatggttccaacaagattattttataacttttataac |
| oCH24 | ggaaccattagaaacagcatagcaagttaaaataaggctagtc |
| oCH25 | gctgttttcattgaatggttccaacaagattattttataacttttataac |
| oCH26 | gttggaaccattcaatgaaaacagcatagcaagttaaaataaggctagtc |
| oCH27 | ccaacaagattattttataacttttataacaaataatcaaggagaaattca |
| oCH28 | gaaaaaaacagcatagcaagttaaaataaggctag |
| oCH29 | aacttgtcatgctgttttcattgaatggttccaacaagattattttataacttttataac |
| oCH30 | tgaaaacagcatgacaagttaaaataaggctagtccgttatcaac |
| oCH31 | cttgctatgctgttttttcaaaacagcatagctctaaaaccaacaagattattttataa |
| oHK55 | ggactagccttattttaacttg |
| oHK156 | cagcatagcaagttaaaataaggc |
| oRAH56 | gtaatacaggggcttttcaagactgaag |
| oRAH59 | ccctctttctcaagttatcatcggc |
| oRAH61 | gactcttcaagtcgatgaaagaaactatcatc |
| oRAH62 | gtttctttcatcgacttgaagagtctttttgg |
| oRAH69 | ccgatgataacgttggatctaaaggtgaagaactgttcac |
| oRAH70 | cctttagatccaacgttatcatcggcaatgttgttttggg |
| oRAH71 | ctctttctcaactatttgtagagctcatccatgccg |
| oRAH72 | gagctctacaaatagttgagaaagagggttaataccagcag |
| oRAH73 | gttcataaataatcatcctcctaaattcatgttttgggaccattcaaaacagc |
| oRAH74 | atgaatttaggaggatgattatttatgaacgttttagagctatgctgttttgaatggtc |
| oRAH75 | ttaaccaagcaatagatgaatttaggaggatgattatttatgaacaggacagactaattaact |
| oRAH76 | agttaattagtctgtcctgttcataaataatcatcctcctaaattcatctattgcttggttaa |
| oRAH81 | atgaatttaggaggatgattattttagaacgttttagagctatgctgttttgaatggtc |
| oRAH82 | gttctaaaataatcatcctcctaaattcatgttttgggaccattcaaaacagc |

| | |
|---|---|
| oRAH83 | ggctcccctatcagcttcaatg |
| oRAH84 | cattgaagctgataggggagcc |
| oRAH86 | ggtcaaatatgttactgaaggaatgcg |
| oRAH97 | gcaaatttagctggtagccctg |
| oRAH167 | ttaaccaagcaatagatgaatttaggaggatgattattttagaacaggacagactaattaact |
| oRAH168 | agttaattagtctgtcctgttctaaaataatcatcctcctaaattcatctattgcttggttaa |
| oRAH298 | ggatttaatttaaactttttattttaggaggcaaaaatg |
| oRAH299 | gcctcctaaaataaaaagtttaaattaaatccataatgag |
| oRAH306 | gggaaaggagttcctgaatttaggatatg |
| oRAH307 | catatcctaaattcaggaactcctttccc |
| oRAH308 | cttaatggaatcggttcatctagtaaggaaaaatta |
| oRAH309 | cttactagatgaaccgattccattaagagg |
| oRAH312 | gcccctgtattactgcatttattaagagtatta |
| oRAH313 | ctcttaataaatgcagtaatacaggggc |
| oRAH318 | tatttttctcgttgttatcatcggcaatgttgttttggg |
| oRAH319 | gccgatgataacaacgagaaaaatataaaacacagtcaaaac |
| oRAH320 | ctataaattatttaataagtaatagttgagaaagagggttaataccagcag |
| oRAH321 | ccctctttctcaactattacttattaaataatttatagctattgaaaagag |
| oRAH384 | ccttattccatcataagattaaaaggaggatgtaataaaatgtcgtggcaccgaattaagcaa |
| oRAH385 | ttgcttaattcggtgccacgacattttattacatcctccttttaatcttatgatggaataagg |
| oRAH387 | gctaacaatgccatgattggtcgg |
| oRAH389 | gttatcatcggcaatgttgttttggg |
| oRAH390 | cccaaaacaacattgccgatgataac |
| oRAH391 | gccgatgataacaacgagaaaaatataaaacac |
| oRAH392 | gtgttttatatttttctcgttgttatcatcggc |
| oRAH393 | gctgttgaataatacagctaacaatgcc |
| oRAH394 | ggcattgttagctgtattattcaacagc |
| oRAH395 | ctttccctacacgacgctcttccgatctNNNNNNNNcttgtatttcttggggaggcttttg |
| oRAH396 | ggagttcagacgtgtgctcttccgatctNNNNNNNNtgagattctaaatctgcatgaatcaattttaaaat |
| oRAH397 | ctttccctacacgacgctcttccgatctNNNNNNNNgatattttaggatttgatgttaactcctcaacc |
| oRAH398 | ggagttcagacgtgtgctcttccgatctNNNNNNNNtcttagtgagatatttgaaaatttgaagtatctctag |
| oRAH399 | ctttccctacacgacgctcttccgatctNNNNNNNNaacgcaaagtgatactatttttgaaaaatgt |
| oRAH400 | ggagttcagacgtgtgctcttccgatctNNNNNNNNgcccctgtattactgcatttattaagagta |
| oRAH407 | catttatttacgttgacgaatcttggagctc |
| oRAH408 | gagctccaagattcgtcaacgtaaataaatg |
| oRAH409 | ggataagattgtttatgaaaatcgaaatcagcc |
| oRAH410 | ggctgatttcgattttcataaacaatcttatcc |
| oRAH411 | gaatcagaaatcatgcttgtaacagatattttagg |
| oRAH412 | cctaaaatatctgttacaagcatgatttctgattc |
| oRAH413 | ctttccctacacgacgctcttccgatctNNNNNNNNatttgagtcagctaggaggtgactg |
| oRAH420 | ggagttcagacgtgtgctcttccgatctNNNNNNNNgcatatcaaacataagtatcattctcatatatctataac |
| oRAH421 | ggagttcagacgtgtgctcttccgatctNNNNNNNNgaatatgacaaggacattctcatccac |
| oRAH422 | ctttccctacacgacgctcttccgatctNNNNNNNNcactatgctggtaaaacggcta |
| oRAH423 | ggagttcagacgtgtgctcttccgatctNNNNNNNNgcatgctcctagatagcaagattg |
| oRAH424 | ctttccctacacgacgctcttccgatctNNNNNNNNcgacgattattgctcaaaagattttgaat |
| oRAH425 | ggagttcagacgtgtgctcttccgatctNNNNNNNNtaaagtataaccataatccagacctgcatt |
| oRAH426 | ctttccctacacgacgctcttccgatctNNNNNNNNtcaagagatttggagcatccaatc |

| | |
|---|---|
| oRAH427 | ggagttcagacgtgtgctcttccgatctNNNNNNNNtttgggaaaaggctgatttcgatt |
| oRAH428 | ctttccctacacgacgctcttccgatctNNNNNNNNggcctttagtggataagattgtttatgaa |
| oRAH429 | ctttccctacacgacgctcttccgatctNNNNNNNNggaaaggagttcctgaatttaggat |
| oRAH430 | ggagttcagacgtgtgctcttccgatctNNNNNNNNtccctcagccgaccaatcat |
| oRAH431 | ctttccctacacgacgctcttccgatctNNNNNNNNctgttgaataatacagctaacaatgcc |
| oRAH432 | ggagttcagacgtgtgctcttccgatctNNNNNNNNggaatcggttcatctagtaaggaaaaa |
| oRAH433 | /5Phos/attaagattcatcaaaagcctcccc |
| oRAH434 | /5Phos/tatttcttaataactaaaaatatggtataatactcttaataaatgcag |
| oRAH454 | ggttcatctagtaatttaaaattaagattcatcaaaagcctcccc |
| oRAH455 | gaatcttaattttaaattactagatgaaccgattccattaagagg |
| oRAH456 | gatttctgatggtttgattgttttcatcttgtgatcaaaaaatttaag |
| oRAH457 | gaaaacaatcaaaccatcagaaatcatgcttgtacagatattttagg |
| oRAH458 | catgcttgtacaagatattttaggatttgatgttaactcctcaac |
| oRAH459 | cctaaaatatcttgtacaagcatgatttctgattctttgattg |
| oRAH460 | gaagtgcagagtttacaagagtatatatcattgacaaatttaacag |
| oRAH461 | ctcttgtaaactctgcacttcatcctttgttagaaaagctcc |
| oRAH462 | ggaagattctaaacttaaatttttgatcacaagatgaaaacaatc |
| oRAH463 | caaaaaatttaagtttagaatcttcctcatattggtaacaatattgc |
| oRAH464 | ggaagattcttatcttaaatttttgatcacaagatgaaaacaatc |
| oRAH465 | caaaaaatttaagataagaatcttcctcatattggtaacaatattgc |
| oRAH466 | gcagatttagaagaacaatttaatgagaaacccgaagtgaaatc |
| oRAH467 | cattaaattgttcttctaaatctgcatgaatcaattttaaaatggttg |
| oRAH468 | gcagatttagaatatcaatttaatgagaaacccgaagtgaaatc |
| oRAH469 | cattaaattgatattctaaatctgcatgaatcaattttaaaatggttg |
| oRAH526 | tcagtcacctcctagctgactc |
| oRAH575 | atcatctgtgttttaaggatgcctataaaacggagctg |
| oRAH576 | catccttaaaacacagatgattattcttataggataatttcgagtggg |
| oRAH577 | caacagctcatctgatgccttttatggtcgtcatg |
| oRAH578 | aggcatcagatgagctgttggtaatcgtttatcatcac |
| oRAH583 | ctaggagcaaactcctattttgaaaaatcccaatctattatgg |
| oRAH584 | caaaataggagtttgctcctagatagcaagattgattc |
| oRAH585 | gctcctatacggaaaaatcccaatctattatggatttatatcatgg |
| oRAH586 | gggatttttccgtataggagcatgctcctagatagc |
| oRAH587 | gagagttatatactttgttttcagatacattttcatataatgg |
| oRAH588 | ctgaaaacaaagtatataactctctctttattttgggaaaagg |
| oRAH589 | ctttgtttaaagatacattttcatataatggtaaagagatgtatc |
| oRAH590 | gaaaatgtatctttaaacaaagtaaataactctctctttattttgggaaaag |
| oRAH591 | cagatacatttatctatataatggtaaagagatgtatctcacgaatattattagc |
| oRAH592 | taccattatagataaatgtatctgaaaacaaagtaaataactctctc |
| oRAH593 | ttttcatatcatggtaaagagatgtatctcacgaatattattagcg |
| oRAH594 | catctctttaccatgatatgaaaatgtatctgaaaacaaagtaaataactc |
| oRAH595 | gagttatagatttatgagaatgatacttatgtttgatatgccg |
| oRAH596 | tcattctcataaatctataactcatatcctaaattcaggaactcc |
| oRAH597 | gatatgccgcaggacaccgctgaggaacg |
| oRAH598 | cagcggtgtcctgcggcatatcaaacataagtatcattctc |
| oRAH599 | ggaacgaatggcctatcgaaaatttcggaaatttttac |
| oRAH600 | ttcgataggccattcgttcctcagcggtgtc |

| | |
|---|---|
| oRAH601 | gcctatcgacaatttcggaaatttttacttagtgaaggg |
| oRAH602 | ttccgaaattgtcgataggcttttcgttcctc |
| oRAH603 | gccatgattactcggctgagggagcataatc |
| oRAH604 | ctcagccgagtaatcatggcattgttagctgtattattc |
| oRAH605 | gcataatcctaaaaaaggaaatattacattactaacggtcacgg |
| oRAH606 | tatttcctttttaggattatgctccctcagcc |
| oRAH607 | agaaataatcatattgcaaactccgatgaaagacttg |
| oRAH608 | gagtttgcaatatgattatttctttcaccatgtaaataaatcattcg |
| oRAH609 | caaactccactgaaagacttgtatttcttggggag |
| oRAH610 | caagtctttcagtggagtttgcaatacaattatttctttcac |
| oRAH611 | gatccggctgctaacaaagcc |
| oRAH612 | atttcgcgggatcgagatctcg |
| oRAH634 | cgagatctcgatcccgcgaaat |
| oRAH679 | ggtaaagagatggctctcacgaatattattagcgattatac |
| oRAH680 | cgtgagagccatctctttaccattatatgaaaatgtatctg |
| oRAH681 | cgatgaagcacttgtatttcttggggaggcttttg |
| oRAH682 | gaaatacaagtgcttcatcggagtttgcaatacaattatttctttcac |
| oRAH683 | ggtaaagagatgcgtctcacgaatattattagcgattatac |
| oRAH684 | cgtgagacgcatctctttaccattatatgaaaatgtatctg |
| oRAH685 | ccgatgaataccttgtatttcttggggaggcttttg |
| oRAH686 | gaaatacaaggtattcatcggagtttgcaatacaattatttctttcac |
| oRAH687 | ggtaaagagatgtttctcacgaatattattagcgattatac |
| oRAH688 | cgtgagaaacatctctttaccattatatgaaaatgtatctg |
| oRAH689 | ccgatgaaaaacttgtatttcttggggaggcttttg |
| oRAH690 | gaaatacaagtttttcatcggagtttgcaatacaattatttctttcac |
| oRAH691 | gttgggctactgttgtggtaaatacccactcg |
| oRAH692 | cacaacagtagcccaaccagccatcagtcacc |
| oRAH693 | tggttggaaaactgttgtggtaaatacccactcg |
| oRAH694 | ccacaacagttttccaaccagccatcagtcacc |
| oRAH695 | gtattgcaaacgccgatgaaagacttgtatttcttggg |
| oRAH696 | ttcatcggcgtttgcaatacaattatttctttcaccatg |
| oRAH697 | ggcttaaagccgctaatcagtttaatcagttcaattttgc |
| oRAH698 | tgattagcggctttaagcccaaactgagtcatacatcc |
| oRAH699 | gatattatggcaccatttaggcctttagtggataagattg |
| oRAH700 | cctaaatggtgccataatatcgctagcaaaattgaactg |
| oRAH701 | gttagcagccggatctcatatcctaaattcaggaactcctttcc |
| oRAH702 | ggatccaccaatctgttctctgtg |
| oRAH703 | aacagattggtggatccatggctggttggcgtactg |
| oRAH704 | gttagcagccggatcttaagattcatcaaaagcctccccaag |
| oRAH705 | aacagattggtggatccatgagttatagatatatgagaatgatacttatgtttgatatgcc |
| oRAH706 | aacagattggtggatccatggctggttgggctactg |
| oRAH707 | aacagattggtggatccatggctggttggaaaactgttg |
